# Development and validation of a streamlined workflow for proteomic analysis of proteins and post-translational modifications from dried blood

**DOI:** 10.1101/2025.09.26.678912

**Authors:** Matthew W. Foster, Youwei Chen, Marlene Violette, Michael T. Forrester, J. Scott Mellors, Brett S. Phinney, Robert S. Plumb, J. Will Thompson, Timothy J. McMahon

**Affiliations:** Division of Pulmonary, Allergy and Critical Care Medicine, Department of Medicine, Duke University School of Medicine, Durham NC USA; Proteomics and Metabolomics Core Facility, Duke University School of Medicine, Durham NC USA; Move Analytical, Carrboro NC USA; Proteomics Core, University of California Davis, Davis CA USA; Waters Corporation, Milford MA USA; VA Medical Center, Durham NC USA

**Keywords:** Mitra device, VAMS, microflow liquid chromatography, capillary zone electrophoresis, data-independent acquisition, HILIC, IMAC, stepped collision energy, Orbitrap Astral

## Abstract

It is increasingly recognized that the ‘omic analysis of whole blood has applications for precision medicine and disease phenotyping. Despite this realization, whole blood is generally viewed as a challenging analytical matrix in comparison to plasma or serum. Moreover, proteomic analyses of whole blood proteomics have almost exclusively focused on (non)targeted analyses of protein abundances and much less on post-translational modifications (PTMs). Here, we developed a streamlined workflow for processing twenty microliters of venous blood collected by volumetric absorptive microsampling that incorporates serial trypsinization, N-glycopeptide and phosphopeptide enrichment and avoids laborious sample dry-down or cleanup steps. Up to 10,000 analytes (reported as protein groups, glycopeptidoforms and phosphosites) were quantified by liquid chromatography-tandem mass spectrometry (LC-MS/MS) in approximately 2 h of MS acquisition time. Using these methods, we explored the stability of “dried” and “wet” blood proteomes, as well as effects of ex vivo inflammatory stimulus or phosphatase inhibition. Multi-omics factor analysis enabled facile identification of analytes that contributed to inter-individual variability of the blood proteomes, including N-glycopeptides that distinguish immunoglobulin heavy constant alpha 2 allotypes. Collectively, our results help to establish feasibility and best practices for the integrated MS-based quantification of proteins and PTMs from dried blood.

## INTRODUCTION

While the “blood proteome” is common nomenclature [1–4], the term is very often a misnomer that instead refers to the cell-free plasma or serum proteome. Whole blood—typically captured as dried blood spots (DBS)—is compatible with both immunoassays and (non)targeted mass spectrometry [5, 6] yet is far less utilized than plasma or serum as a starting matrix for proteomic studies. Because whole blood is easily sampled and preserved at point-of-collection, it may be less susceptible to pre-analytical variability as compared to blood plasma, which can be confounded by hemolysis, clotting or platelet activation [7, 8]. Dried blood has been used extensively for small molecule analysis in newborn and genetic screening, as well as drug testing, but proteomic applications are being described with more frequency [9–11].

Volumetric microsampling (VAMS) has emerged as an alternative to collection of DBS. Neoteryx Mitra devices contain a hydrophilic porous material designed to collect 10-30 µL blood, and are—in contrast to DBS—reported to be insensitive to hematocrit (i.e. red blood cell concentration) while offering a convenient format for remote sampling, specimen storage and processing [12, 13]. Mitra devices are compatible with typical sample processing workflows for mass spectrometry-based and affinity proteomics. Proof-of-concept studies have assessed “on-tip” analyte stability and assay precision, studied the variability due to sampling source (e.g. capillary versus venous), modeled diet-induced proteome changes and explored the application to health surveillance [13–18].

The enormous dynamic range of protein abundances in blood-based biofluids presents unique analytical challenges. State-of-the-art approaches designed to increase depth of proteome coverage—namely “next-generation” affinity proteomics, nanoparticle corona and extracellular vesicle enrichment [19–21]—have been applied to dried blood in a few instances [22, 23]. However, equal or greater focus has been applied to methods that use differential protein precipitation or extraction from dried blood specimens, such as sodium carbonate precipitation [10] or lithium chloride extraction [24]. In contrast to protein enrichment or depletion, methods aimed at increasing the breadth of the blood proteome, rather than maximizing depth, might also be advantageous for discovery-based ‘omics.

Recently, Shen et al. reported a multi-omic approach for lipidomics, metabolomics and proteomics from a single Mitra device, and applied this method to “metabolic” and “physiome” profiling [15]. Nonetheless, there are numerous classes of analytes that remain to be explored, including abundant post-translational modifications (PTMs). Protein N-glycosylation and phosphorylation are among the most abundant mammalian PTMs, and in blood, they serve as important modulators of immunity and inflammation [25–28]. There are only a few descriptions of N-glycopeptide analysis in whole or dried blood [29, 30], but none that has comprehensively profiled the N-glycoproteome. Similarly, phosphoproteomes have been enriched from tryptic digests of erythrocytes [31, 32], but not, to our knowledge, from whole blood. Here, we sought to develop and validate methods for serial enrichment coupled to mass spectrometry-based analysis, of N-glycopeptides and phosphopeptides from dried blood that could be easily integrated into a robust bottom-up proteomic workflow.

## MATERIALS AND METHODS (see Supplementary Material for additional methods)

### Blood collection

Whole blood was collected from normal human subjects using Duke IRB Protocol Pro00041793 and collected in BD Vacutainer Li-Heparin or EDTA tubes followed by mixing according to manufacturer’s protocol.

### Ex vivo blood treatment

Freshly collected heparinized blood was transferred to Eppendorf Lobind tubes and treated with or without 1000 ng/mL *E. coli O111:B4)* lipopolysaccharide at 37 °C for 6, 24 or 48 h. At 6 h, blood was also treated with 1 mM sodium orthovandate, 500 nM calyculin A and 50 µM deltamethrin for 1 h. At each endpoint, blood was mixed and loaded onto 20 µL Mitra devices, followed by transfer of clamshell packages to a sealed foil bag containing dessicant packages and drying for 2 h at room temperature (room temp) before storage at -80 °C. Plasma was prepared by centrifugation of blood at 2700 *xg*, 10 min at 4 °C, and the supernatant was collected and stored at -80 °C.

### Wet blood stability study

Freshly collected EDTA-blood was loaded onto Mitra devices, then dried for 2 h at room temp followed by transfer to septum-capped 750 µL Matrix tubes (ThermoFisher). Remaining blood was transferred to 1.5 mL Protein Lobind tubes and incubated at room temp for 2 or 4 h; or 4 °C for 2, 4 or 16 h before loading onto Mitra devices and drying as described above.

### Dried blood stability study

Freshly collected EDTA-blood was loaded onto Mitra devices and clamshells were stored at -80 °C or in a Secador 1.0 Desiccator Cabinet (Belart) containing ∼100 g Drierite at 4 °C or room temp, with relative humidity of ∼10%, for 1d, 3d or 15 d. Tips were transferred to Matrix tubes and stored at -80 °C.

### Sample processing for nontargeted proteomics

290 microliters of 5.5% w/v sodium deoxycholate (SDC) in 50 mM ammonium bicarbonate (AmBic) containing 10 mM dithiothreitol (DTT) was added to a Matrix tube containing a Mitra tip followed by heating at 80 °C for 25 min on a Thermomixer (Eppendorf) fitted with a 500 µL DWP adapter and heated lid at 850 rpm. After cooling, samples were alkylated with 15 µL of 400 mM iodoacetamide (IAM) in AmBic at room temp in the dark for 30 min. IAM was quenched by addition of 15 µL of 200 mM DTT in AmBic, and digestions were performed by adding 20 µL of 12.5 mg/mL TPCK-treated bovine trypsin (Worthington) in AmBic followed by incubation at 37 °C for 2 h on a Thermomixer. Reactions were quenched by addition of 38 µL of 20/20/60 (v/v/v) TFA/MeCN/water to each sample (note: pipette tip should not touch the DOC solution) followed by vortexing to fully precipitate the SDC. After a brief centrifugation, samples were transferred to an Isolute Filter+ plate (Biotage) using a multichannel pipettor and wide-bore pipette tip, and filtered using a positive pressure manifold (Biotage Pressure+). The filtrate was captured in 750 µL Matrix tubes. A study pool QC (SPQC) sample was made by mixing equal volumes of all samples.

Plasma was processed as above with the following modifications. An SPQC sample was made prior to digestion and was processed in three independent replicates. Twenty microliters of plasma was reduced by adding 200 µL of 5.5% SDC and 11 mM DTT in AmBic followed by reduction at 80 °C and alkylation with 20 µL of 250 mM IAM and subsequent quenching with 10 µL of 200 mM DTT. Digestion used 20 µL of 5 mg/mL TPCK trypsin followed by quenching with 30 µL of 15/20/66 v/v/v TFA/MeCN/water.

### LC-MS/MS of blood and plasma proteome analysis

Tryptic digests were analyzed by LC-MS/MS using a Waters ACQUITY UPLC interfaced to a ThermoFisher Exploris 480 mass spectrometer. Analyses of Mitra samples used 3-5 µL of peptide digests, and analyses of plasma samples used 10 µL of peptide digests, respectively. For autosampler injection, tubes were sealed with a Storage Mat III (Costar). After direct injection, peptides were separated on a 1 x 100 mm or 1 x 150 mm ACQUITY Premier 1.7 µm CSH C18 column (Waters) using a flow rate of 100 µL/min, a column temperature of 55 °C and a gradient of 0.1% (v/v) formic acid (FA) in H_2_O (MPA) and 0.1% (v/v) FA in MeCN (MPB) as follows: 0-60 min, 3-28% MPB; 60-60.5 min, 28-90% MPB; 60.5-62.5 min, 90% MPB; 62.5-63 min, 90-3% MPB; and 63-67 min, 3% MPB. A PEEK tee (ThermoFisher cat# P-727) was used post-column to introduce a solution of 50% (v/v) dimethyl sulfoxide (DMSO) in MeCN at 6 µL per min. In addition, for analysis of whole blood, a solvent divert was used to deliver 50/49.9/0.1 (v/v/v) MeCN/H_2_O/FA to the source, and flow from the analytical column to waste, during the first 2 min and last 5 min of the gradient as previously described [33].The LC was interfaced to the MS via a Optamax NG ion source using heated electrospray ionization (HESI) with the following tune parameters: sheath gas, 32; aux gas, 5; spray voltage, 3.5 kV; capillary temperature, 275 °C; aux gas heater temp, 125 °C.

MS analysis used staggered-overlapping data-independent acquisition (DIA) [34–36]. Briefly, a MS1 scan used a 120,000 resolution from 375-1600 m/z, AGC target of 300% and maximum injection time (IT) of 45 ms, and MS/MS was performed using targeted MS2 (tMS2) with 30,000 resolution, automatic gain control (AGC) target of 1000%, a maximum IT of 60 ms, and a NCE of 30. Data was collected in centroid mode. The tMS2 method used an inclusion list generated in EncylopeDIA [37] with 28 m/z windows and 22 windows per loop. Gas phase fractionation was performed on six replicates of the SPQC pool using DIA acquisition as described above except that each DIA method used 18 windows, an 8.5 m/z isolation width and optimized window placement.

### Identification and quantification of unenriched blood and plasma

Raw MS data was converted to *.htrms format using HTRMS converter (Biognosys) and processed in Spectronaut 19 (Biognosys). A spectral library was generated from DIA data using Spectronaut Pulsar. Searches used a UniProt database with *Homo sapiens* specificity (downloaded on 04/22/24) and appended with sequences for yeast ADH1, bovine cationic and porcine trypsin, lysyl endopeptidase, a concatemer of previously identified variant peptides and additional entries for ApoL1 C-terminal variants (20,434 total entries) [36]. Default settings were used (e.g. Trypsin/P specificity; up to 2 missed cleavages; peptide length from 7-52 amino acids) with the following modifications: fixed carbidomethyl(Cys), variable protein N-terminal acetylation (blood only) and semitryptic N-terminal specificity.

Library matching and quantification in Spectronaut used default extraction, calibration, identification and protein inference settings. Quantification was performed at MS2 level using q-value setting, local normalization [38] and MaxLFQ [39] protein roll-up.

### N-glycopeptide enrichment

Dual enrichment hydrophilic interaction chromatography (HILIC) enrichment with Ti-IV-IMAC (**Supplemental Methods**) [40], or conventional HILIC, were performed with modifications. Briefly, a 10% (w/v) slurry of Polyhydroxyethyl A (12 µm, 300 Å; PolyLC cat# BMHY1203) in 0.1% TFA was incubated for 30 min. A spin tip containing three discs of MK360 filters (Ahlstrom cat# 3600-0470) and 200 µL slurry was fabricated as described in **Fig. S1** and washed with 2 x 200 µL of 80% (v/v) MeCN:H_2_O containing 0.1% TFA (“HILIC wash buffer”). Eighty microliters of tryptic digest was adjusted with 325 µL of a 320:4 (v/v) mixture of MeCN and TFA, for a final of 80% (v/v) MeCN and 1% (v/v) TFA. Approximately 200 µL of the diluted digest was loaded onto the spin tip followed by centrifugation, and this was repeated for the remaining volume. Sample loading was repeated twice more followed by washing with 3 x 200 µL HILIC wash buffer and elution with 2 x 200 µL of 0.1% TFA. Eluents were lyophilized and reconstituted in 30 µL of 0.1% FA. Alternatively, 40 µL of eluents were diluted with 160 µL 0.1% FA and loaded directly onto Evotips.

### LC-MS/MS of N-glycopeptide-enriched samples

LC-MS/MS used a Waters M-class LC in trap-elute configuration (**Supplemental Methods**) or Evosep One LC. The Evosep One LC used a 60 sample-per-day method, Bruker Pepsep 8 cm x 150 μm column (1.5 μm particle size), PepSep Sprayer and 30 μm stainless steel emitter. The LC was interfaced to a ThermoFisher Exploris 480 and Orbitrap Astral MS using a Nanospray Flex Source. Stepped-collision energy data-dependent MS/MS (SCE-DDA) used the Exploris 480 MS with a 60,000 resolution precursor mass range from 650-2000 m/z, 300% AGC and 25 ms IT, and a 30,000 resolution MS/MS scan, with a 200% AGC and auto IT, an isolation window of 1.2 m/z and stepped NCE of 20, 30 and 40%. Cycle time was 1 s.

Normalized collision energy data-independent acquisition (glyco-DIA) used the Orbitrap Astral MS with an Orbitrap precursor scan every 0.6 s at 240K resolution from 850-1550 m/z, 300% AGC, 50 ms IT. DIA MS/MS in the Astral analyzer used 139 x 5 m/z windows from 850- 1550 m/z with a 500% AGC, 8 ms IT and 35% NCE.

#### Identification and quantification of N-glycopeptides

N-glycopeptides were identified and quantified from SCE-DDA data using GlycoDecipher 1.05 and GlyPep-Quant v.1.00 [41] with default parameters except for semitrypsin specificity, up to 2 missed cleavages and carbamidomethyl(Cys) fixed modification. Quantification with GlyPep-Quant used match between runs selected. The “glycopeptide” output was filtered to remove rows annotated with “glycan stepping” mass shifts [42] and additional steps were performed in R, including quantile normalization, extension of the peptide sequence and removal of peptides lacking a N-X-S/T motif (X is not Pro). The amino acid positions of modified Asn were determined (note: multiple N-X-S/T motifs in the same peptide were retained) and an “Accession” was created with Uniprot protein name, glycosylation site and abbreviated glycan composition. Finally, redundant accessions were numerically labeled, starting with smallest (e.g. semitryptic or no missed cleavage) sequence.

Analysis of Astral glyco-DIA data used the glyco-N-DIA workflow of FragPipe [43–45] followed by visualization in Skyline [46]. Default Fragpipe workflow parameters were used except for semi-N-terminal trypsin specificity, no variable modifications and Human_Nglycans_large-708 database.

### Phosphopeptide enrichment

Phosphopeptide enrichments used TiO spin tips [47] or Ti-IMAC magnetic particles [48] as previously described with some modifications. For TiO spin-tip enrichments, 80 µL of tryptic digests (or HILIC unbound fractions) were adjusted with solid glutamic acid (∼30 mg), 225 µL MeCN and 6 µL TFA (65% v/v MeCN and 2% v/v TFA with saturated GA; Glu binding buffer), or with a solution containing 30 mg glycolic acid, 320 µL MeCN and 4 µL TFA (80% v/v MeCN, 1% v/v TFA and 1 M GA; GA binding buffer). Enrichments were performed using 3 mg TiO spin-tips (GL Sciences). After pre-elution and equilibration with binding buffer, samples were loaded at 1,000 x*g* for 5 min, and loading was repeated. Washes and elutions used 100 µL at 1000 x*g* for 2min. For Glu excluder, tips were washed twice with Glu binding buffer, followed by 65% MeCN/0.5% TFA and 5% MeCN/0.1% TFA. For GA excluder, tips were washed twice with GA binding buffer followed by 80% MeCN/1% TFA, and 20% MeCN/0.1% TFA. Both methods used sequential elutions of 5% aq. NH_4_OH and 5% aq. NH_4_OH with 50% MeCN. Eluents were adjusted to pH <6 with neat formic acid and lyophilized.

For Ti-IMAC enrichment, 300 µL of HILIC flow-through was adjusted with 100 µL of 20% TFA (v/v) in MeCN and 1 M glycolic acid (GA) followed by enrichment using a Kingfisher Flex (ThermoFisher) using 40 µL of MagReSyn® Ti-IMAC HP (Resyn Biosciences). Enrichment used the manufacturer’s protocol except that 0.25 M GA was utilized in binding and wash buffers.

Peptides were eluted with 200 µL of 1% NH_4_OH followed by acidification with 50 µL of 10% formic acid, and up to 100 µL was diluted with 150 µL of 0.1% formic acid before loading onto Evotips.

#### LC-MS/MS of phosphopeptide-enriched samples

LC-MS/MS used an Evosep One at 60SPD interfaced to a ThermoFisher Exploris 480 or Orbitrap Astral as described above. MS/MS on the Exploris 480 used a 60,000 resolution precursor scan from 380-950 m/z with an AGC target of 1000% and max IT of 60 ms, and a staggered/overlapping DIA method using the tMS2 scan function with 16 x 32 m/z windows covering a mass range of 400-928 m/z with 30,000 resolution, AGC of 1000% and 60 ms IT. MS/MS on the Orbitrap Astral used a 240,000 resolution precursor scan from 380-1080 m/z with 500% AGC and 50 ms IT with a cycle time of 0.6 s. DIA used 139 x 5 m/z windows from 380-1080 m/z with a 500% AG, 8 ms IT and 28% NCE. Additional analyses using ZipChip CE-MS/MS or Evosep timsTOF MS/MS as described in **Supplemental Methods**.

#### Phosphopeptide identification and quantification

DIA data was analyzed in Spectronaut 19 after (with demultiplexing of staggered/overlapping Orbitrap DIA data) and conversion to .htrms format. Pulsar searches used variable N-terminal protein acetylation and phospho(S/T/Y) with a maximum of 25 fragment ions and minimum of 3 fragment ions. Identification and quantification were performed in Spectronaut at a 1% FDR and local normalization was applied on phosphorylated precursors. For quantification, normalized precursor abundances were exported with PTM localization set to 0, and the data was filtered to remove precursors that were not localized (0.75 probability cutoff) in at least 10% of samples. Missing values were not imputed unless otherwise noted. For the phosphatase inhibitor datasets, data were not normalized, and background imputation was performed in Spectronaut.

### Statistical analysis

Analyses were performed in R (coding aided by ChatGPT-4o) or ChatGPT- imbedded Python, Graphpad Prism, Clustvis [49] or ggVolcanoR server [50]. Multi-omics factor analysis used the MOFA2 R-package [51] and model training with 6 factors. Additional analyses used k-nearest neighbor imputation (k=5) and log2 transformation prior to statistical analyses.

Comparisons between groups used paired t-tests with Benjamini-Hochberg FDR correction. Fold change calculations were the average of paired ratios for each individual.

#### Figure design

Final figures were made in Inkscape. Some illustrations used Biorender via an academic license to Duke University Department of Medicine.

## RESULTS

### Proteomic analysis of Mitra devices using sodium deoxycholate-assisted digestion and microflowLC-MS/MS

We first sought to develop and validate a facile method for processing of a 20 µL Mitra device (∼5 mg blood) that would be compatible with downstream PTM enrichment. We have previously utilized deoxycholate-assisted trypsin digestion for targeted proteomics of whole blood [33] and we adapted this approach for processing of Mitra devices (Fig. 1A). The end of the device (“Mitra tip”) containing dried blood was easily dislodged when sealing a Matrix tube with a septum cap, thus providing a convenient solution for sample storage and processing [52]. Sample heating and mixing steps were performed in a deep-well plate adapter to minimize plate-based “edge effects” (**Fig. S2A**) [53]. Following acid precipitation (using a repeater pipette to avoid touching the pipette tip to SDC solution during addition), samples were transferred to a filter plate using a wide bore tip. Since the Mitra tip remained at the bottom of the Matrix tube, it was challenging to transfer the entire sample to the filter plate. Still, we were able to consistently recover >200 µL of digest (∼2.5 mg assuming quantitative recovery).

**Figure 1.**
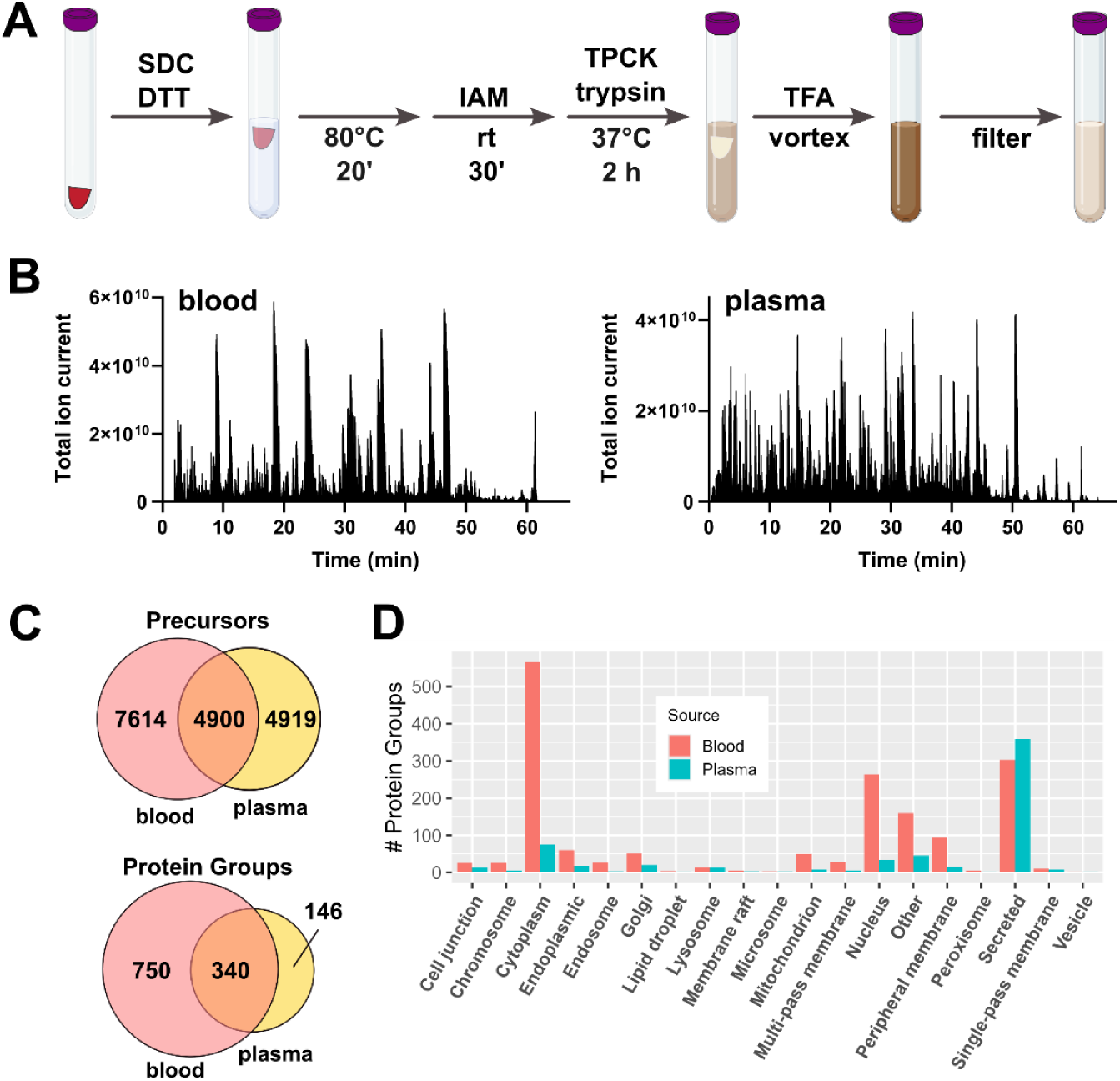
Deoxycholate-assisted digestion and non-targeted proteomic analysis of dried blood compared to matched plasma. **A**) Processing steps for sodium deoxycholate (SDC)-assisted trypsin digestion of 20 µL whole blood stored on Mitra devices. Abbreviations are defined in Methods. Plasma processing is nearly identical and was as described previously [36]. **B**) Total ion current (TIC) for matched venous blood (∼0.125 µL) and plasma (∼0.67 µL) digests analyzed by microflowLC-MS/MS with post-column addition of DMSO. **C**) Venn diagrams of precursors and protein groups quantified in blood and plasma. **D**) Number of Uniprot cellular localization terms among protein groups quantified in blood versus plasma.

The blood proteome was analyzed using direct injection of 5 µL digest (0.125 µL blood or 60 µgs; Fig. 1B) directly from Matrix tubes (**Fig. S2B**), separation by 1 mm scale microflow LC (1 x 100 mm), a 100 µL/min flow rate and a 60 min gradient [35, 54, 55], with post-column addition of DMSO [56]. To minimize source contamination absent a cleanup step, we used a divert valve to waste the first 2 min and last 5 min of a 67 min method [33]. MS data were collected on an Orbitrap Exploris 480 using staggered/overlapping data-independent acquisition (DIA) [34]. For comparison, 20 µL of plasma from the same n=3 controls was digested in a final volume of 300 µL, and 10 µL of digest (0.67 µL plasma or 33 µgs) was analyzed (Fig. 1B). Using direct-DIA for peptide/protein identification, there were ∼12,500 precursors and 1,090 protein groups quantified in the three blood samples versus ∼9,800 precursors and 486 protein groups in plasma. There were approximately 50% more precursors uniquely quantified, but ∼5x more protein groups uniquely quantified, in blood versus plasma (Fig. 1C). As expected, the blood proteome was enriched in cellular compartment and organelle-localized proteins but had similar number of secreted proteins as compared to plasma (Fig. 1D).

We further evaluated gas phase fractionation to populate the spectral library [34]. In a representative analysis of samples from four healthy control donors, addition of six GPF samples increased identified protein groups in the spectral library by ∼50% to >2,000 but increased total quantified proteins by only ∼25% to 1,519 (**Supplemental Data**). The GPF-enhanced library resulted in slightly lower precision compared to the direct-DIA of individual samples alone (median CV of 4.4% versus 3.6%, respectively), as measured by replicates of a study pool quality control (SPQC) sample, made by combining equal volumes of all samples post-digestion.

### Effects of inflammatory stimulation of the blood versus plasma proteomes

Having established a workflow for non-targeted proteomic analysis of whole blood, we wanted to further explore differences in plasma and blood in an *ex vivo* model (Fig. 2A). Treatment of whole blood with *E. coli* lipopolysaccharide (LPS) has been used to assess cell type-specific chemokine expression [57] and monocyte function [58]. Typically, blood is diluted with cell culture medium prior to stimulation and the readout (e.g. tumor necrosis factor-alpha quantification) is performed on the cell-free fraction (i.e. plasma) [58]. We incubated non-diluted lithium-heparin venous blood (shown to yield higher cell activation compared to EDTA-blood [59]) obtained from three normal donors with or without 1000 ng/mL LPS for 6, 24 and 48 h. After gently resuspending cells by inverting the incubation tubes, blood was loaded on Mitra devices, and plasma was collected after centrifugation. Following trypsinization, LC-MS/MS was performed on block-randomized samples in singlicate and interspersed with at least three replicates of an SPQC sample. As in our pilot studies (Fig. 1C), approximately twice as many protein groups were quantified in blood (1,180) versus plasma (630), with similar precision (median %CV ∼3.8 and 4 for blood and plasma, respectively) based on n=3 analyses of the SPQC pools.

**Figure 2.**
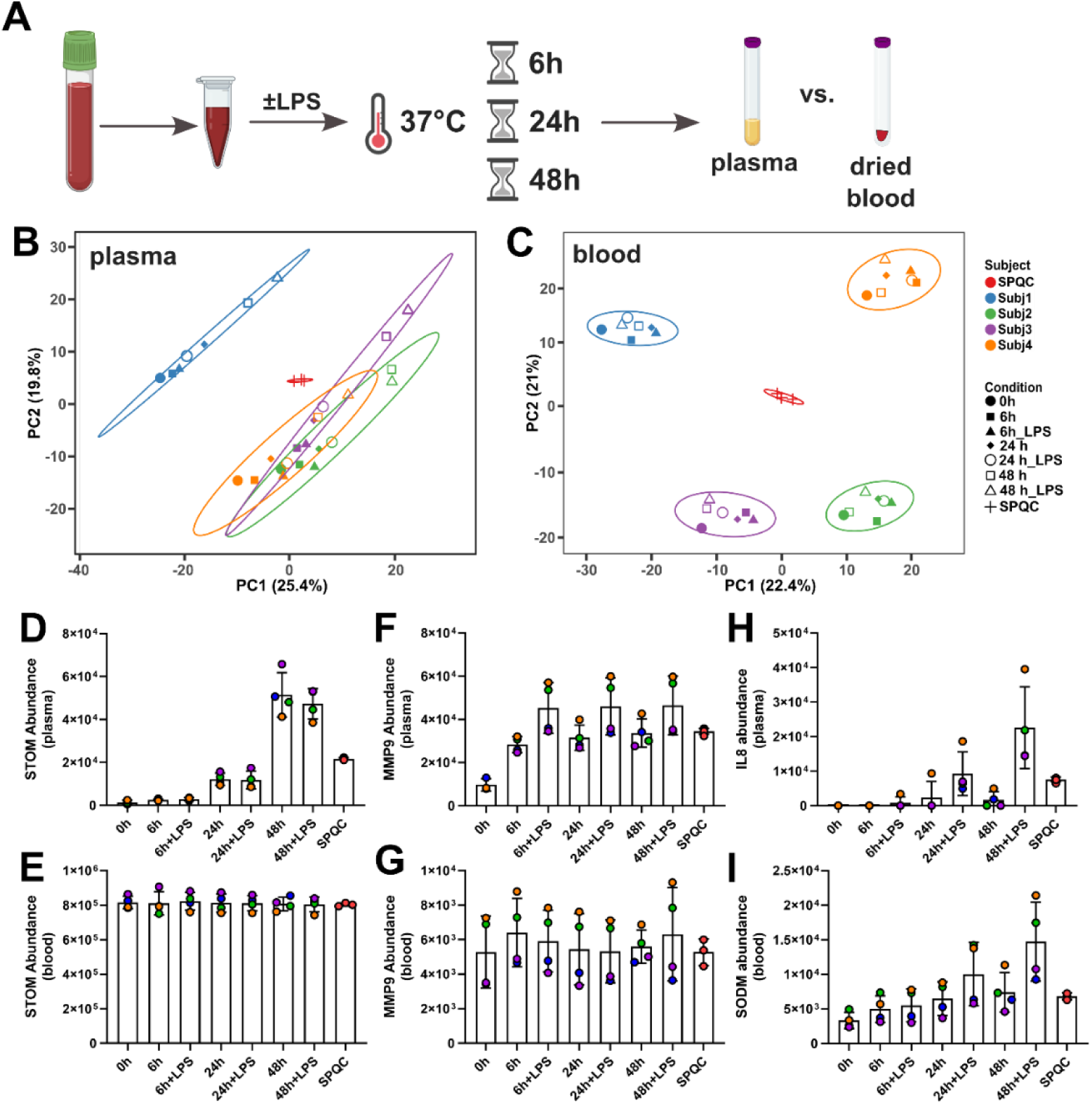
Plasma and whole blood proteome changes with ex vivo inflammatory stimulus. **A**) Heparinized blood from four healthy control donors (Subjects 1-4) was loaded onto Mitra tips or centrifuged to recover plasma after blood draw (0h) or treatment ± 1000 ng/ml *E. coli* lipopolysaccharide at 37 °C for 6, 24 or 48 h followed by proteomic analysis. SPQC samples were n=3 prep and n=2 technical replicates (plasma) and n=3 technical replicates (blood). **B-C**) Principal component analysis of (**B**) plasma proteome and (**C**) blood proteome. **D-I**) Abundances of: stomatin (STOM) in (**D**) plasma and (**E**) blood; matrix metalloproteinase 9 (MMP9) in (**F**) plasma and (**G**) blood; **H**) interleukin-8 (IL8) in plasma; and **I**) mitochondrial superoxide dismutase (SODM). Donors in **D-I** are colored as in **B-C**. Line and error bars are mean ± s.d, n=4 per group.

Principal component analysis (PCA) of plasma samples showed shifting of samples along the positive PC1/PC2 diagonal (Fig. 2B) whereas the blood samples more tightly clustered by donor (Fig. 2C). A time-dependent increase in hemoglobins and many other erythrocyte proteins, including the integral membrane protein stomatin (Fig. 2D), was observed in plasma but not in whole blood, which was insensitive to changes in protein compartmentalization (Fig. 2E). Although statistical significance was not reached due to variability to individual responses (**Supplemental Data**), neutrophil-derived proteins, including lactotransferrin, matrix metalloproteinase 9 (MMP9; Fig. 2F), neutrophil elastase, azurocidin and myeloperoxidase were also increased in plasma (but not whole blood) after 6 h of incubation, suggestive of leukocyte degranulation.

The relative change in plasma was proportional to the total abundance of these proteins in whole blood (Fig. 2G). For example, subjects 2 and 4 had the largest increase in MMP9 in LPS-stimulated plasma as well as the highest total amounts of MMP9 in blood at all timepoints. The inflammatory cytokine interleukin-8 was increased in plasma at 24 and 48 h of LPS treatment (Fig. 2H) but not detected in blood, whereas mitochondrial superoxide dismutase was one of the few proteins that was specifically increased in blood with LPS exposure (Fig. 2I). These data begin to reveal how the treatment of whole blood can have modest influence on the blood proteome but significant effects on the plasma proteome.

### Enrichment of N-glycopeptides from whole blood digests

For ease of throughput and scalability, we sought to develop PTM enrichment methods that would be compatible with the SDC digest without an intermediate solid phase extraction. Given the large dynamic range in blood, we decided to utilize 80 µL of tryptic digests (∼1 mg based on estimated concentration of 250 mg/mL). This allowed us to dilute the digests to starting conditions for downstream enrichments (e.g. up to 80% MeCN) while still maintaining volumes suitable for 96-well formats. Since we viewed these studies as highly exploratory, we decided to prioritize throughput over coverage, focusing on relatively short (∼30 min or less) LC-MS/MS analyses.

We first tested dual enrichment of N-glycopeptides and phosphopeptides using CAE- Ti(IV)-IMAC (Fig. 3A) [40]. Eighty microliters of digest pooled from healthy controls were adjusted to 80% MeCN and 3% TFA in a final volume of 400 µL and loaded onto 20 mg CAE-Ti-IMAC packed into P200 pipette tips (**Fig. S1**). Elutions used decreasing MeCN/0.1% FA (for glycopeptides) or 10% NH_4_OH (for phosphopeptides). After lyophilization, fractions were analyzed by nanoLC-MS/MS using stepped collision energy DDA [60], an empirically-optimized precursor range of 650-2000 m/z, and Orbitrap MS/MS. Putative phosphopeptide-enriched fractions were analyzed by staggered/overlapping DIA. Using Glycounter [61] to assess both oxonium ion-containing MS/MS scans and likely glycopeptide MS/MS spectra, the first fraction (60% MeCN/0.1% FA) exhibited highest enrichment compared to 40% MeCN and 20%-0 MeCN (Fig. 3B); however, N-glycopeptide peptide spectral matches returned by Glyco-Decipher [41] dropped precipitously after the 60% MeCN fraction (Fig. 3B). Further, the basic elution (10% NH_4_OH) with stepped MeCN from 60-0% yielded essentially no identifiable phosphopeptides (data not shown). Even though the CAE-Ti-IMAC resin did enrich for N-glycopeptides, it was expensive (∼$1000/g or ∼$20/sample), and elution with 60% MeCN required a subsequent dry-down step prior to LC-MS/MS. Thus, we sought an alternative approach. We instead tested a HILIC method using spin tips packed with 20 mg of Polyhydroxyethyl A (PolyA) resin (∼$30/g or ∼$0.6 per sample; Fig. 3A) [62]. Digests were adjusted to 80% MeCN and 1% TFA for loading, and elution was performed with 0.1% TFA. This approach also yielded a high percentage of oxonium ion-containing MS/MS (Fig. 3C), and a larger number of identified glycopeptides per sample compared to CAE-Ti-IMAC (Fig. 3D). Liquid chromatography using the Evosep One exhibited better MS utilization compared to nanoLC (**Fig. S3A**), and the PolyA HILIC method could be directly loaded onto Evotips without additional cleanup or drying steps, simplifying the workflow and avoiding sample loss (**Fig. S3B**). The vast majority of identified N-glycopeptides, and the top 20 based on number of site-specific glycans, were from soluble/secreted proteins (Fig. 3F). Two major erythrocyte membrane glycosites that were identified included Asn45 of glucose transporter 1 (Glut1/SLC2A1) and Asn642 of band 3 anion transporter (B3AT/SLC4A1; **Supplemental Data**). The most abundant glycan was the di-sialylated Hex(5)HexNac(4)NeuAc(2) (Fig. 3G).

**Figure 3.**
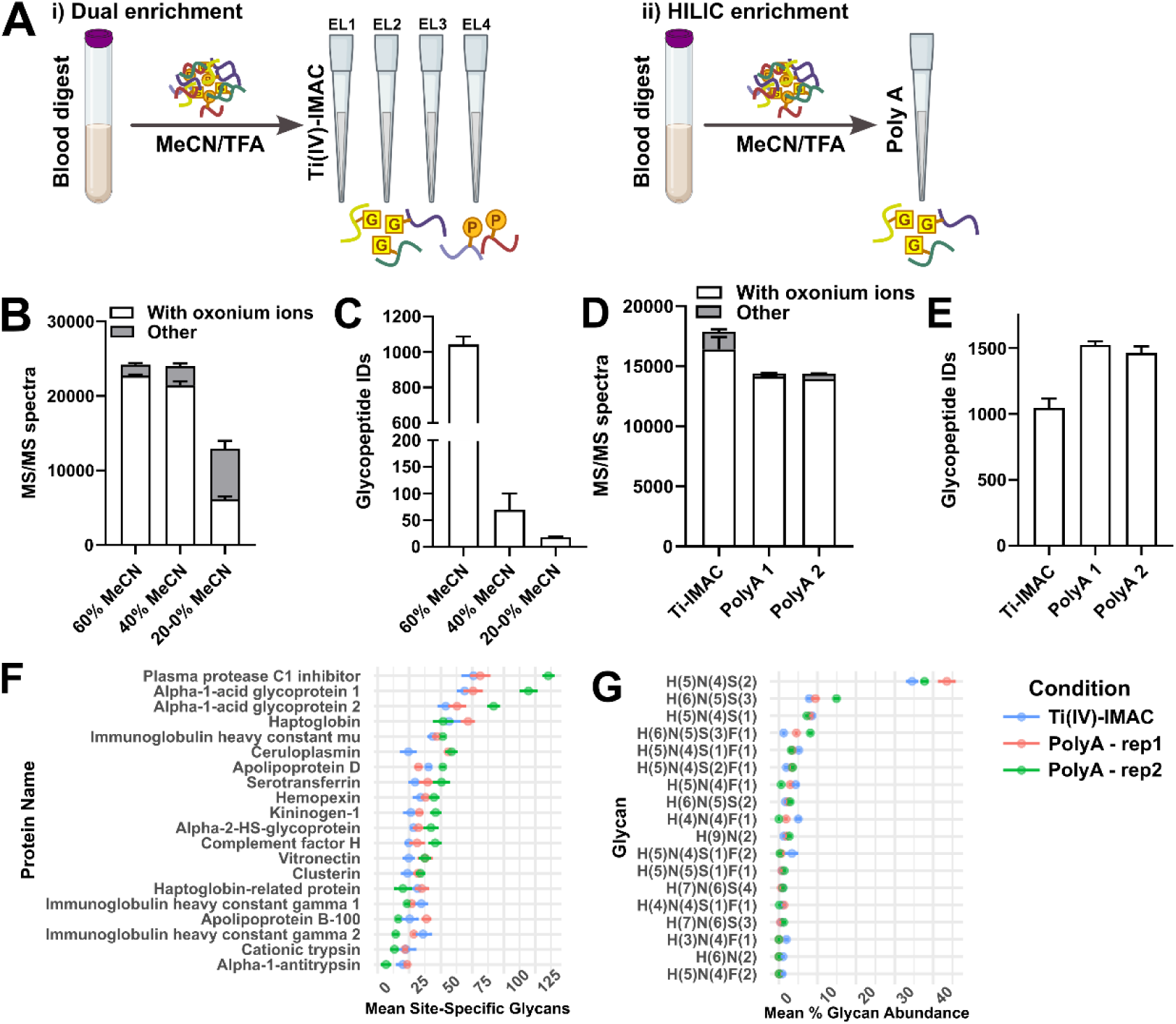
N-glycopeptide enrichment from blood using immobilized metal affinity chromatography and hydrophilic interaction chromatography. **A**) Summary of approaches evaluated for glycopeptide enrichment. Tryptic digests of Mitra tips were pooled from normal human controls and enriched for N-glycopeptides (**see Methods**) using (**i**) CAE-Ti(IV)-IMAC (“Ti-IMAC”) or (**ii**) Polyhydroxyethyl A (“PolyA”) spin tips. Ten percent of eluents were analyzed using a 15 min nanoLC gradient (Ti-IMAC) or 60 SPD Evosep LC method (PolyA) and SCE-DDA. **B-C**) Data from stepwise Ti-IMAC eluents were analyzed using GlyCounter (in **B**) or GlycoDecipher (in **C**). (**D-G**) Data from Ti-IMAC eluents (60% MeCN/0.1% FA) and two independent PolyA eluents (0.1% FA) were analyzed by Glycounter (in **D**) and Glyco-Decipher (**E-G**). Data are mean ± s.d (n=2-3 replicates pre group).

### Glycopeptide quantification using data-independent acquisition (DIA)

Data-independent acquisition has been increasingly utilized for glycoproteomics, although the data processing workflows are still limited. Nonetheless, we viewed DIA as a potentially useful method for validation of differentially-expressed peptides, as both MS1 and MS2 chromatograms can be used to establish identity and for relative quantification. As proof-of-concept for glyco-DIA analysis of the blood proteome, we used Orbitrap Astral DIA-MS/MS, which has been recently explored for N-glycopeptide quantification [63]. For Glyco-DIA analyses, we used 5 m/z windows from 850- 1550 m/z to include the majority of the precursors identified in DDA analyses and to insure sufficient points-per-peak. The method also took advantage of parallelization to collect precursor scans every 0.6 s at 240,000 resolution, with a DIA duty cycle of 1.5 s as compared to a 60,000 precursor resolution and 1 s cycle time for the Orbitrap SCE-DDA method. We reanalyzed the wet blood dataset as proof-of-concept followed by data processing using the “glyco-N-DIA” workflow in FragPipe v23, with generation of a spectral library from SCE-DDA data, quantification in DIA-NN and visualization in Skyline (**Supplemental Data**). These data were used for validation of differential glycopeptide abundances identified in multiomic factor analysis (MOFA; see below).

### Parallel and serial enrichment methods for phosphopeptides from whole blood

Failing to detect phosphopeptides from the Ti(IV)-IMAC eluents, we next tested parallel phosphopeptide enrichment with titanium oxide (TiO) spin tips (Fig. 4A, i), adjusting 80 µL of digest with MeCN, TFA and saturating glutamic or 1 M glycolic acid (Glu or GA) as excluders to desired concentrations. The Glu protocol (adapted from Li et al. [47]) utilized 65% MeCN, resulting in a lower starting volume for enrichment, but Glu needed to be added as a solid to reach a saturated solution. The Glu protocol yielded about twice as many quantified precursors (1820 versus 990) and higher specificity (68% versus 43%, respectively) as compared to GA using sample-specific spectral libraries (Fig. 4B), with similar trends in the abundance of phosphorylated and non-phosphorylated precursors (Fig. 4B). As with the N-glycopeptide analysis, we adapted this workflow to the Evosep One LC with a 60SPD method, which shortened the inject-to-inject time by ∼50%. However, we did include a drying step prior to sample loading, as the phosphopeptide elution protocols from TiO used up to 50% MeCN.

**Figure 4.**
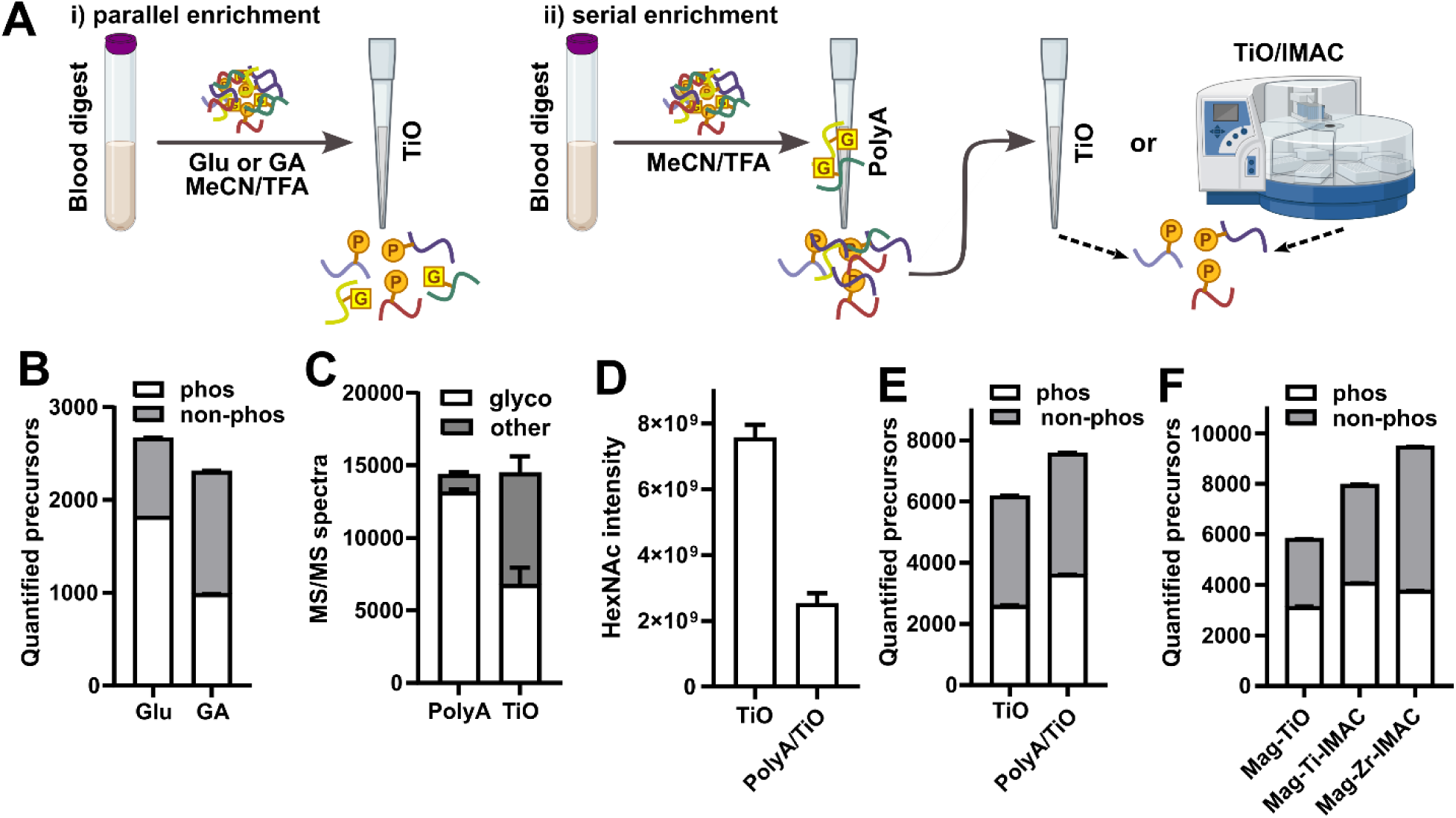
Methods for phosphopeptide enrichment from whole blood digests. **A**) Summary of approaches evaluated for phosphopeptide enrichment. **B-F**) 80 µL of tryptic digests (∼1 mg) were diluted to binding conditions and enriched for post-translational modifications followed by analysis of ∼40% of phospho-enriched or ∼10% of glyco-enriched fraction by LC-MS/MS. **B**) Glutamic acid (Glu) or glycolic acid (GA) were tested as excluders using TiO spin tips and analyzed by OT-DIA. **C**) Glycopeptide MS/MS were predicted from SCE-DDA analysis of Polyhydroxyethyl A or TiO-enriched fractions using Glycounter. **D**) HexNAc (204.087 m/z) oxonium ion was quantified using the “extract ion” option in DIA-NN. **E-F**) Phosphopeptides were quantified in Astral-DIA analysis of (**E**) TiO versus serial PolyA/TiO enriched samples and (**F**) magnetic nanoparticle-enriched samples. Data in **F** are n=2 per condition, and others are n=3 per condition. Spectronaut was used for phosphopeptide identification and quantification. Note spectral libraries were generated, and analyses were performed, on each sample group independently.

Despite the feasibility of the parallel TiO enrichment, we observed a large number of oxonium ion-containing MS/MS spectra in the TiO-enriched sample. Indeed, when TiO-enriched samples were analyzed by SCE-DDA, ∼50% of the peptide spectral matches (PSMs) were predicted to be glycopeptides (Fig. 4C). This raised the idea that the unbound flow-through from the PolyA spin tip could be utilized for serial enrichment of phosphopeptides (Fig. 4A**, ii**). Neat tryptic digests, or PolyA/HILIC flow-throughs, were adjusted to TiO-binding conditions followed by phosphopeptide enrichment and Orbitrap Astral DIA. Serial enrichment resulted in a >100-fold reduction in HexNAc (204.087 m/z) intensity in phosphopeptide-enriched samples, as assessed by oxonium ion scanning [64] (Fig. 4D), as well as a larger number of quantified phosphopeptide precursors as compared to TiO enrichment without glycopeptide depletion (Fig. 4E).

Finally, we also tested the compatibility of magnetic nanoparticle-based phosphopeptide enrichment using the serial enrichment workflow (Fig. 4A**, ii**). Unbound flow-throughs from PolyA/HILIC spin tips were adjusted to loading conditions with either 1 M glycolic acid (for TiO nanoparticles) or 0.25 M GA (for Ti-IMAC and Zr-IMAC nanoparticles). Eluents (1% NH_4_OH) were acidified and loaded directly onto EvoTips. The number and abundance of localized phosphopeptide precursors was higher with Ti-IMAC and Zr-IMAC magnetic nanoparticles as compared to TiO nanoparticles (Fig. 4F). We chose the Ti-IMAC nanoparticles for further quantitative evaluation based in part on a separate test of non-human mammalian blood (unpublished), although there might be advantages of Zr-IMAC because of its longer reported shelf-life. Serial enrichments with PolyA and either TiO or Ti-IMAC were utilized in wet blood stability studies (see below).

Phosphorylation within the red blood cell is important for signal transduction and deformability [65] and has been studied in both isolated RBCs and purified membranes [32, 66], but not (to our knowledge) using whole blood directly. As a proof-of-concept, we enriched phosphopeptides from whole blood following treatment with LPS (Fig. 2A) and a cocktail of Ser/Thr and Tyr phosphatase inhibitors (calyculin, deltamethrin and sodium orthovandate [67]) at 6 h post blood draw (**Fig. S4A**). There were 1,542 phosphosites (1,120 pSer, 338 pThr and 83 pTyr) quantified in blood from n=4 healthy controls. Similar to the unenriched proteome, LPS had a negligible effect on phosphorylation (**Supplemental Data**) whereas PI treatment resulted in a significant increase (avg. log2FC >1; q<0.1) in 303 phosphosites (**Fig. S4B**). Numerous RBC membrane proteins, including erythrocyte membrane protein band 4.1 (EBP41), spectrin beta chain (SPTB1) and ankyrin-1 (ANK1; **Fig. S4C**) were among those with the highest number of differentially abundant phosphosites. While only a subset of phosphosites in these proteins were affected, all eleven quantified phosphosites in alpha synuclein (SYUA) were increased with phosphatase inhibition (**Fig. S4B-C**). The majority of circulating SYUA (>99%) is localized to the erythrocyte [68], and these eleven phosphosites—including positional isomers that were distinguishable by retention time (**Supplemental Data**)—are far more than have been previously described in RBCs. Phospho-SYUA in RBCs has been suggested as a biomarker of Parkinson’s disease and other neurodegenerative diseases (multiple system atrophy and progressive supranuclear palsy)[69–72]. These data suggest that mass spectrometry may have utility for phospho-synuclein quantification in blood and establish a workflow that should be compatible with targeted quantification.

We further tested the compatibility of this approach with LC-MS/MS using an Evosep One LC and Tims-TOF with DIA-PASEF acquisition, as well as solid phase extraction capillary zone electrophoresis (SPE-CZE)[73] coupled to Orbitrap DDA MS/MS. Similar results were observed between the three analyses (**Fig. S4D and Supplemental Data**). SPE-CZE-DDA-MS/MS appeared to favor the detection of larger peptides, which could be a factor of the trapping conditions, the greater mass range spanned by DDA-MS/MS versus DIA-MS/MS and the separation properties of CZE. We do also acknowledge the limitation of LFQ-DDA-MS/MS for the assignment of positional isoforms, motivating translation of this approach to DIA [73].

### Assessing stability of proteins and PTMs in “dried” and “wet” blood

Given the potential use of whole blood for remote sampling and clinical proteomics, there have been a number of studies of proteome stability during sample collection, transport and storage [6, 15, 74, 75]. To extend this to PTMs, and to test the entirety of sample processing and analysis workflows, we separately modeled short-term storage of “dried” and “wet” venous EDTA-blood from four healthy controls per study. In the dried blood stability study (n=28 samples excluding pools), Mitra devices were loaded with blood, dried and either immediately transferred to -80 °C (i.e. t0) or incubated at 4 °C or room temp for 1, 3, 7 or 15 days prior to storage at -80 °C (Fig. 5A). In the wet stability study (n=24 samples excluding pools), a t0 sample was collected, or whole blood was stored at 4 °C for 2, 4 and 24 h, and at room temp for 2 and 4 h, before capturing on Mitra devices (Fig. 5B). The dried blood samples were processed and analyzed earlier in the method development process, and included parallel PolyA/HILIC and TiO enrichments, while wet blood samples used serial enrichment using both TiO and Ti-IMAC and included Orbitrap Astral DIA analysis of both phosphopeptides and N-glycopeptides (Fig. 5C).

**Figure 5.**
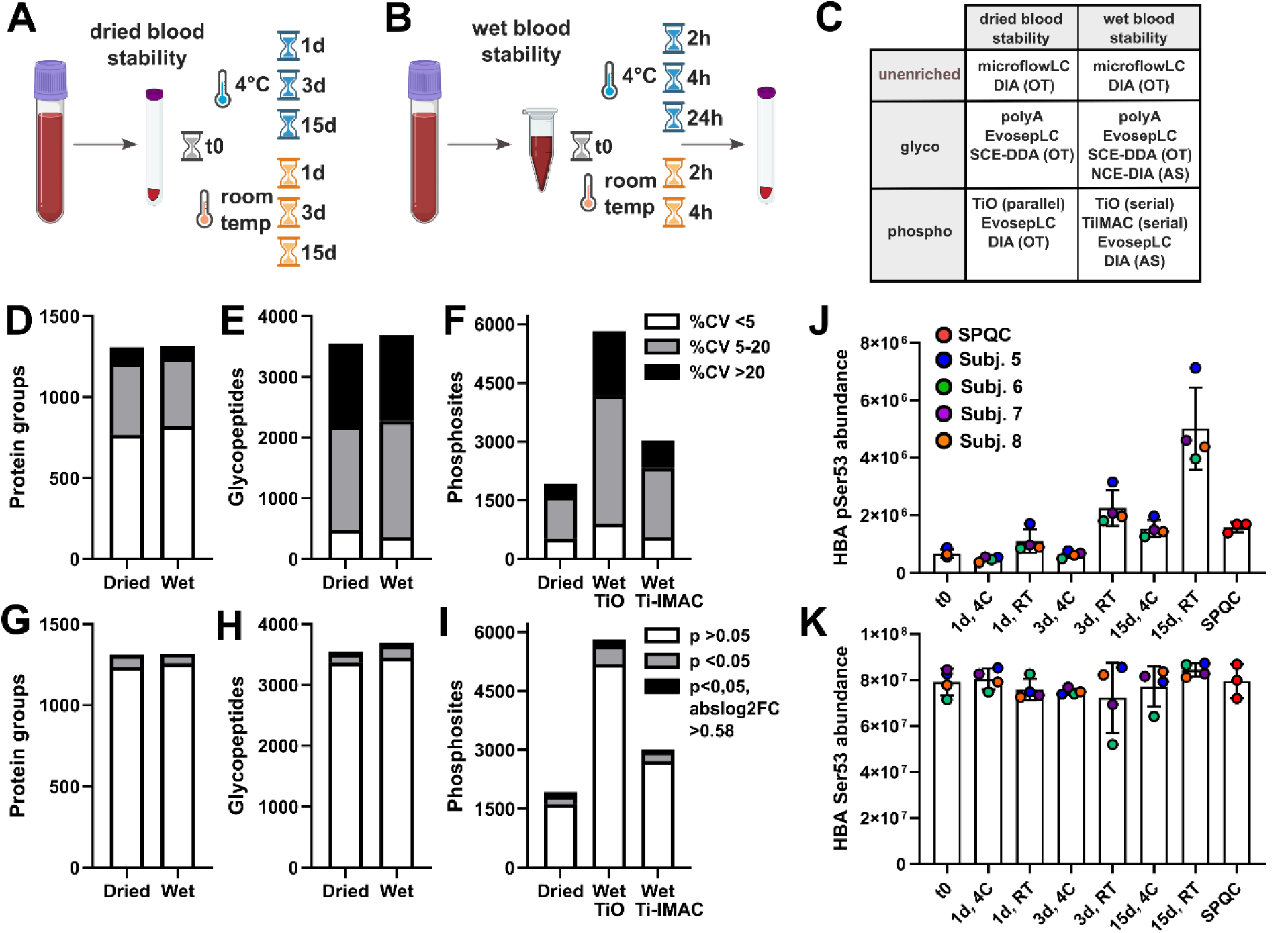
Modeling effects of dried and wet blood storage on the whole blood proteome. Venous EDTA whole blood from four healthy controls per experiment was collected on Mitra tips and transferred to -80 °C after drying for 2 hours (“t0”), and (**A**) stored in a desiccator at 4 °C and room temp for 1,3 and 15 days or (**B**) incubated at 4 °C and room temp for 2 and 4 h, or 24 h at 4 °C prior to loading on Mitra tips. **C**) Summary of enrichment and LC-MS/MS methods applied to the two stability studies. **D-F**) Percent coefficients of variation were determined for n=3 technical replicates of (**D**) post-digestion and (**E-F**) post-enrichment SPQC samples for dried and wet blood stability studies. (**G-I**) Summaries of numbers of significant changes in protein and PTM abundance (by paired t-test) between t0 versus 15 d at room temp (dried blood stability) or t0 versus 24 h at 4 °C (wet blood stability). **J**) Storage time-dependent changes in hemoglobin alpha chain (HBA) pSer53 phosphorylation observed in dried blood. **J**) Individual HBA pSer53 phosphosites and **K**) HBA precursor (TYFPHFDLSHGSAQVK3+) from unenriched data visualized across the four donors and SPQC samples.

We filtered datasets to remove missing rows from technical replicates of the SPQC samples. There were 1,308 protein groups, 3,547 glycopepeptides and 1,925 phosphosites quantified in the dried blood stability study. Similarly, the wet blood stability study had 1,317 quantified protein groups, 3,688 quantified glycopepeptides and 5,827 or 3,024 quantified phosphosites with TiO and Ti-IMAC enrichment, respectively (Fig. 5C). Using the technical replicates of the SPQC samples to assess precision, the majority of protein groups quantified had median CVs of <5%, and the majority of quantified PTMs had median CVs of <20% (Fig. 5D**-F**). Statistical testing between the most extreme conditions (e.g. dried blood, 15 d at room temp versus t0; or wet blood (24 h at 4 °C versus t0) was used as a preliminary assessment of analyte stability (**Supplemental Data and** Fig. 5G**-I**). Only 13 analytes passed an FDR-corrected p<0.1 and log2FC>0.58 at the extremes of storage conditions; all of these were phosphosites in the dried blood stability study, with pSer53 of hemoglobin alpha chain (HBA) exhibiting a marked increase in abundance upon room temp storage (Fig. 5J **and Supplemental Data**) with no change in the total amount of Ser53 peptide quantified in the unenriched proteome (Fig. 5K).

### Multi-omics factor analysis across matched blood proteome and PTM-enriched samples

We utilized the “multi-omics factor analysis” R-package (MOFA2)[51, 76] to further explore the dried and wet blood stability datasets (Fig. 6A). This approach allowed easy exploration across proteomes and was robust to missing data. When visualized by a latent factors plot of factors 1 and 2, samples clustered tightly by subject, and SPQCs were centered at zero on x- and y-axes (Fig. 6B and 6D), further demonstrating both the overall precision of the approach and the stability of dried and wet blood under simulated storage conditions.

**Figure 6.**
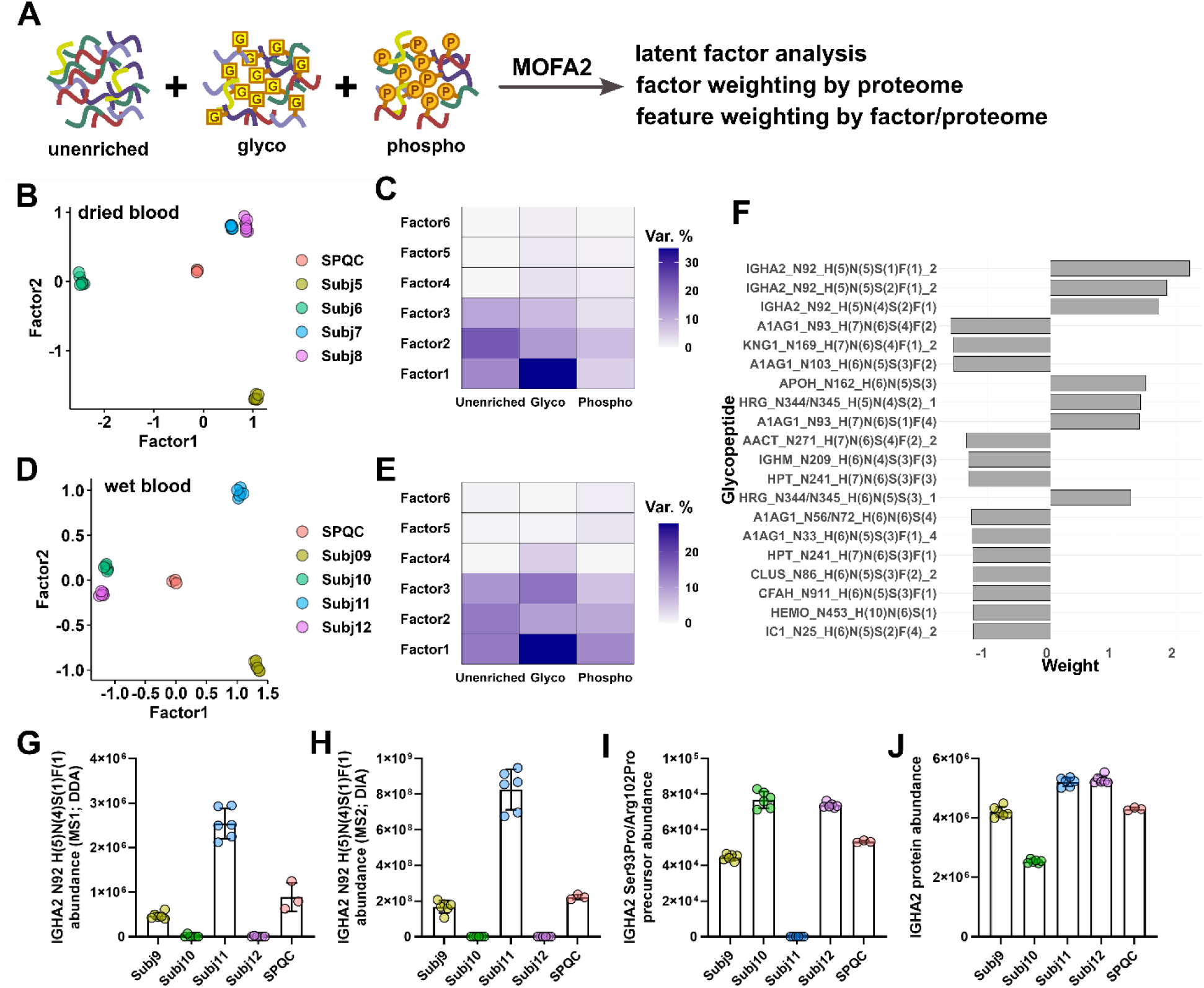
Multi-omics factor analysis (MOFA) of dried and wet blood proteomes. **A**) Unenriched, N-glyco- and phospho-proteomes from dried and wet blood stability studies were analyzed using the MOFA2 R-package. **B and D**) Latent factor plots of factors 1 versus 2 for dried and wet blood stability studies, respectively. **C and E**) Percent variance explained by each proteome within factors 1-6 for dried and wet blood stability studies, respectively. **F**) Top 20 N-glycopeptides weighting factor 1 of the wet blood stability proteome labeled by protein name, modification site, abbreviated glycan composition (H = Hexose, N = HexNAc, S = NeuAc and F = Fucose) and replicate number (for sites represented by more than one peptide due to missed cleavage or semi-specific cleavage). **G-H**) Abundances of Asn92 glycopeptide of IGHA2 with H5N4S1F1 composition in wet blood dataset acquired by (**G**) SCE-DDA and (**H**) NCE-DIA. **I-J**) Abundance of (**I**) IGHA2 precursor (HYTNPSQDVTVPCPVPPPPPCCHPR.3) with Ser93Pro and Arg102Pro substitutions and (**J**) IGHA2 protein in unenriched proteome of wet blood stability dataset. Data in **B** are n=7 per subject and n=3 for SPQC samples, and data in **D** and **G-J** are n=6 per subject and n=3 for SPQC samples.

In both datasets, the N-glycoproteome explained more of the variance of factor 1 versus the unenriched- and phospho-proteomes (Fig. 6C and 6E), and in the wet blood dataset, three immunoglobulin heavy constant alpha 2 (IGHA2) Asn92 (HYTN*SSQDVTVPCR) glycopeptides were the most highly weighted in factor 1 of the N-glycoproteome (Fig. 6G). The A2m(2) allotype of IGHA2 is glycosylated at Ser93, whereas the A2m(1) allotype has a Ser93-to-Pro mutation that eliminates the N-X-S/T (X ≠ Pro) motif, as well as an Arg102Pro substitution [77]. In the SCE-DDA dataset (Fig. 6F), and in an Orbitrap Astral glyco-DIA validation dataset (Fig. 6G **and Supplemental Data**), the A2m(2) glycopeptide was detected in subjects 9 and 11 but not in subjects 10 and 12. The resulting non-glycosylated A2m(1) peptide HYTNPSQDVTVPCPVPPPPPCCHPR was also measured in the unenriched data and was not detected in subject 11 (Fig. 6H). These data suggested that subject 9 was heterozygous A2m(1)/A2m(2), subjects 10 and 12 were homozygous A2m(1) and subject 11 was homozygous A2m(2). IGHA2 protein abundances did not correlate with assigned allotype (Fig. 6I).

MOFA was also useful for extracting the proteins and phosphosites that separated healthy control proteomes (**Supplemental Data and Fig. S5**). To cross-validate the phospho-enrichment approaches, we performed MOFA on the wet blood stability study using phosphosite data from either TiO or Ti-IMAC enrichment. Among the top twenty phosphosites weighting factor 1, seven were shared between the two approaches (**Fig. S5A-B**). Phosphothreonine 2288 of spectrin alpha chain, erythrocytic 1 (SPTA1), phosphoserine 135 of GTP-binding nuclear protein Ran and phosphoserine 267 of copper chaperone for superoxide dismutase (CCS) had a consistent pattern of expression with TiO (**Fig. S5C-E**) or Ti-IMAC enrichment (**Fig. S5F-H**). While the relative abundances of these sites were up to ∼3-fold different between subjects, the relative levels of SPTA1, RAN and CCS (**Fig. S5I-K**) were mostly unchanged. Phosphorylation of Ser135 of Ran increases during mitosis [78, 79], suggesting that this site may be a marker of relative mitotic activity in blood mononuclear cells.

## DISCUSSION

The “blood proteome” is primarily derived through analyses of plasma or serum [8, 20, 80]. However, whole blood can be easier to sample and exhibits lower pre-analytical variability compared to plasma [81, 82], and a greater number of proteins can be quantified from neat blood versus plasma. Compared to serum, blood suffers from the same dynamic range problem and can present a more analytically challenging matrix for proteomics. Improved depth-of-coverage in dried blood has been addressed using affinity proteomics [83–85] and by depletion of high abundance proteins [10, 24]. Here, we developed a relatively simple and scalable approach for blood proteomics that utilizes orthogonal enrichment of abundant circulating PTMs (Fig. 7). Across healthy controls, blood proteomes are mostly affected by inter-individual variability rather than sample storage conditions, and unique information is reflected in each of the measured analyte classes. The assay is relatively easy to scale to a 96-well format—and throughput and coverage of the unenriched proteome can be improved by combining microflow LC with Orbitrap Astral DIA—as recently demonstrated for a longitudinal sepsis cohort, and we further improved throughput and coverage of the unenriched proteome by combining microflow LC with Orbitrap Astral DIA [52]. Each of the three blood proteomes revealed unique information about sepsis resolution, including reduction of markers of acute phase response, inflammation and neutrophil activation. Collectively, these data demonstrate that this “multi-proteomic” approach (i.e. integrated protein and PTM analyses), which utilizes only 20 µL of blood, is both tractable for repository-scale studies and can be useful for phenotyping disease.

The serial enrichment of N-glycopeptides and phosphopeptides is enabled by shared binding conditions between HILIC and IMAC (e.g. 80% MeCN and 1-5% TFA). In addition to the CAE-Ti(IV)-IMAC material that we tested [40], Li and coworkers have also described the synthesis of an epoxy-ATP-Ti(IV)-IMAC resin and utilized it for dual enrichment from mammalian cells and tissues, with an improvement in enrichment specificity for phosphopeptides [86]. Sequential Fe-IMAC and HILIC enrichment (“TIMAHAC”) [87], and integrated IMAC-HILIC spin tips [88], have been developed by Chen et al. and used to enrich N-glycopeptides and phosphopeptides from suspension trap (S-trap) digests of plant tissue. Further, the recently described SCASP-PTM method utilizes serial TiO and Ti-IMAC enrichment downstream of antibody-based enrichment for quantification of up to four PTM classes in cells and tissues [89]. A single tip enrichment of N-glycopeptides and phosphopeptides from blood using an approach like the IMAC-HILIC method might still be an obtainable goal and would simplify large-scale studies. We also recognize the potential for improving the specificity of phosphopeptide enrichment from whole blood, as ∼80% has been claimed for dual enrichment from tissues or cell digests (albeit with much lower %enrichment of N-glycopeptides) [87]. Nonetheless, the findings described here serve as an important starting point for future method refinements.

Despite the extensive adoption of plasma proteomics for population-based studies [90], there are many applications where dried (whole) blood “multi-proteomics” should excel (in addition to sepsis phenotyping, which we have recently demonstrated [52]). While our methods are optimized for Mitra devices, they are not exclusive to a single dried blood platform. Collection of dried blood enables easy remote and microsampling, and DBS standard-of-care in newborn screening. Affinity proteomics has been used to profile herpes simplex virus infection [84] and to predict risk for type 1 diabetes [85] from newborn DBS. Dried blood multi-proteomics should be readily translatable from DBS and open up additional opportunities to profile health and disease in newborns. The blood multi-proteome may also reveal new molecular details of red blood cell dysfunction in hemoglobinopathies, including sickle cell trait and sickle cell disease—that are not captured by plasma proteomics [36, 91]—and may be useful in following response to newly approved gene therapies. Finally, minimally invasive microsampling of dried blood has advantages in studies of non-human and non-model organisms and should benefit both laboratory- and field-based research. Due to the paucity of affinity reagents, mass spectrometry-based approaches are already the norm for plasma and serum proteomics in non-model organisms [92]. Transition to dried blood should be easily adopted for both sample collection and MS-based analysis, thus providing unique opportunities to study the circulating proteome in the context of human disease.

## ACKNOWLEDGEMENTS

This work was supported by funding from the National Institutes of Health, R33-GM146142 (M.W.F and T.J.M.) and R01-HL161071 to T.J.M.

## SUPPLEMENTAL DATA

Supplemental data and methods accompany this submission. Raw data and associated metadata and results have been deposited to the ProteomeXchange Consortium (PXD068610) via the MassIVE repository (MSV000099243 and ftp://MSV000099243@massive-ftp.ucsd.edu). And Skyline files have been uploaded to Panorama Public: https://panoramaweb.org/4PIjSy.url. Code and associated processed data are available at https://github.com/mwfoster/Mitra_proteomics.

