## Supplemental Material for "Development and validation of a streamlined workflow for proteomic analysis of proteins and post-translational modifications from dried blood"

### SUPPLEMENTAL METHODS

**TI(IV)-IMAC dual N-glycopeptide and phosphopeptide enrichment.** Ti-IV-IMAC enrichment were performed as described [1] with minor modifications. Briefly, 20 mg of CAE-Ti-IMAC (J&K Scientific LLC) was swelled with 200 µL of 0.1% TFA and incubated for 30 min. A spin tip was made as described for Polyhydroxyethyl A HILIC (**Fig. S1**). Two hundred microliters of the slurry was added to each tip. The resin was packed by centrifugation at 300 *xg* for 3 min in a swinging bucket rotor followed by washing with 2 x 200 µL of 80% (v/v) MeCN:$H_2O$ containing 3% TFA (wash buffer). Fifty microliters of tryptic digests were adjusted with 200 µL of MeCN and 7.5 µLTFA (final 80% MeCN and 3% TFA). Approximately 200 µL of was loaded onto the spin tips followed by centrifugation, and this was repeated for the remaining volume. Sample loading was repeated for a total of 3 times. Tips were washed 6 times with 200 µL wash buffer, once with 200 µL of 80% MeCN/0/1% FA and eluted with 200 µL of 60% MeCN/0.1% FA, 40% MeCN/0.1% FA, 20% MecN/0.1% and 0.1% FA. The last two fractions were combined. Phosphopeptides were eluted with 60% MeCN/10% $NH_4OH$, 40% MeCN/10% $NH_4OH$ and 10% $NH_4OH$ and acidified with neat FA. Phosphopeptude eluents were combined. All fractions were lyophilized and reconstituted in 30 µL of 0.1% FA.

**Nanoscale liquid chromatography.** LC-MS/MS used a Waters M-class LC in trap-elute configuration. Briefly, peptides Peptides were trapped on a Symmetry C18 180 µm × 20 mm trapping column (Waters) at 5 µL/min with 99.9/0.1 v/v H2O/MeCN and separated on a 75 µm × 10 cm HSS-T3 analytical column (Waters) at 400 nl/min and 55 °C. A gradient of 5-30% mobile phase B (99.9

**LC-timsTOF-MS/MS analysis of blood phosphopeptides.** Phosphopeptides were enriched using 3 mg TiO spin tips and lyophilized. After reconstitution in 0.1% FA, 25% of eluents were loaded onto Evotips and analyzed using an Evosep One LC and Bruker TimsTOF Pro2 MS. The Evosep used a 60 SPD method and the TimsTOF used DIA-PASEF with 50 Da Windows 300-

1200 Da, no overlap in IM or m/z, 1/KO 0.6-1.5, ramp time of 75 ms and cycle time of 0.57 s. Data was processed in Spectronaut 19 as described in Methods.

**Solid phase extraction capillary zone electrophoresis (SPE-CZE) Orbitrap-MS/MS analysis of blood phosphopeptides.** Phosphopeptides were enriched as described above and diluted 1:10 with 100 mM ammonium acetate, 1% acetonitrile before sample injection. SPE-CZE was as previously described [2] with the following modifications. Sample was loaded onto the SPE bed for 480 s (~ 4 µL total volume). MS/MS utilized an Exploris 480 MS in DDA mode with a 45,000 resolution precursor scan from 400-1600 m/z, AGC of 300% and IT of 50 ms. Up to 5 MS/MS were performed per cycle using an intensity threshold of 5E4, charge state 2-5, 10 s dynamic exclusion, 2 m/z isolation window, 15,000 resolution (from 0-7 min) or 30,000 resolution (from 7-14 min), 200% AGC, auto IT and NCE of 30%. Data was analyzed in Skyline using MS Amanda with variable pSTY with MS1 quantification [3], and data was filtered to include putative phosphatase inhibitor-sensitive peptide precursors.

### SUPPLEMENTAL FIGURES

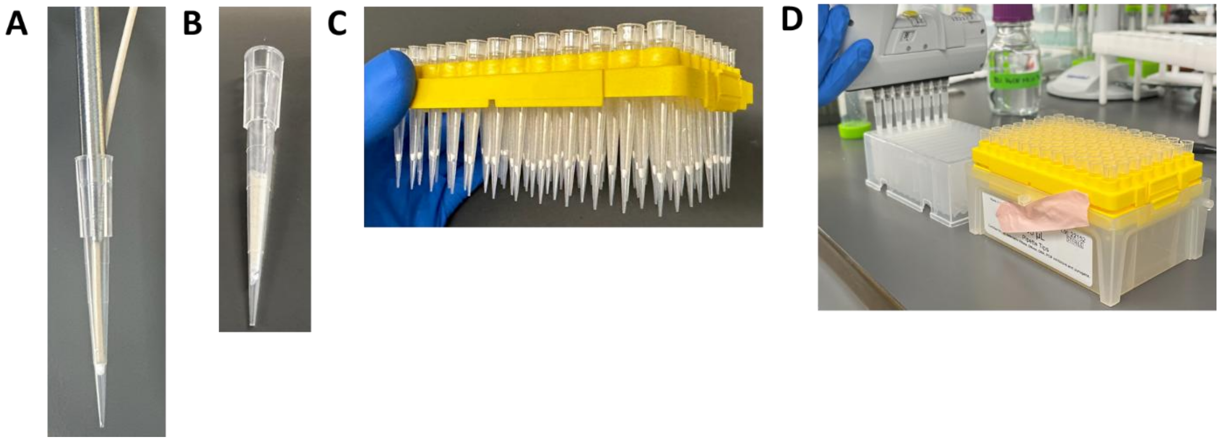


**Figure S1. Fabrication of spin tips for N-glcyopeptide enrichment. A**) A 2 mm leather punch (Jmuiiu Store; Amazon.com) was used to cut three discs of MK360 quartz filters (Ahlstrom cat# 3600-0470), which were packed into a 200 µL pipette tip using 1/16th inch o.d. PEEK tubing. **B**) Two hundred microliters of a 10% w/v solution of pre-swelled CAE-Ti-IMAC or Polyhydroxyethyl A (12 µm, 300 Å; PolyLC cat#

BMHY1203) was added to the tip followed by centrifugation at 200 *xg* for 3 min in a swinging bucket rotor. **C**) An empty peptide tip rack and **D**) multichannel pipettor were used to load up to 96 tips at time.

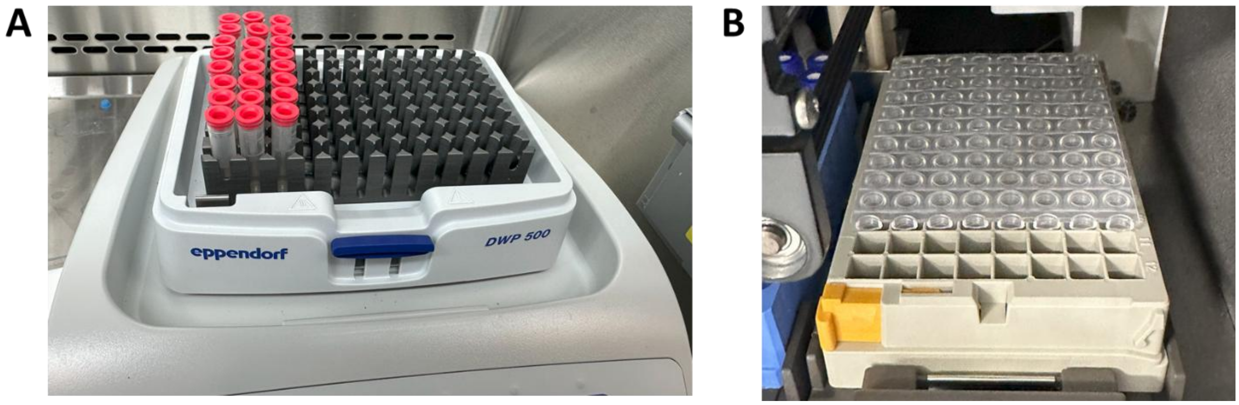


**Figure S2. Figure S2. Sample processing and analysis using Matrix tubes. A**) Samples incubation steps, including reduction and trypsin digestion were performed in a Thermomixer with a deep-well plate (DWP500) and heated lid to minimize edge effects associated with heating using a flat bottom plate adapter. **B**) Following filtration of SDC-precipitated digests into clean Matrix tubes, they could be sealed with a Capmat III (Costar; cut to fit the number of tubes in a batch) and placed in an LC autosampler for analysis of the unenriched proteome.

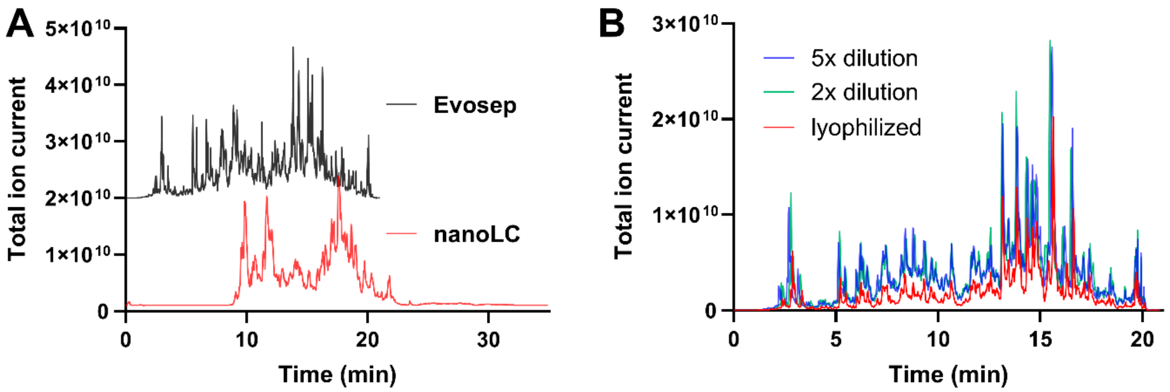


**Figure S3. LC-MS/MS of N-glycopeptide-enriched samples. A**) Representative TIC of an N-glycopeptide-enriched fraction separated using a trap-elute nanoLC with a 75 µm x 10 cm analytical column and 15 min gradient of 5-30% MeCN versus 150 µm x 8 cm analytical column and Evosep LC 60SPD method. **B**) N-glycopeptides were eluted from Polyhydroxyethyl A resin using 0.1% FA and diluted 2- to 5-fold with 0.1% FA before loading onto and Evotip, or lyophilized and reconstituted before loading on an Evotip followed by analysis using a 60SPD Evosep LC method.

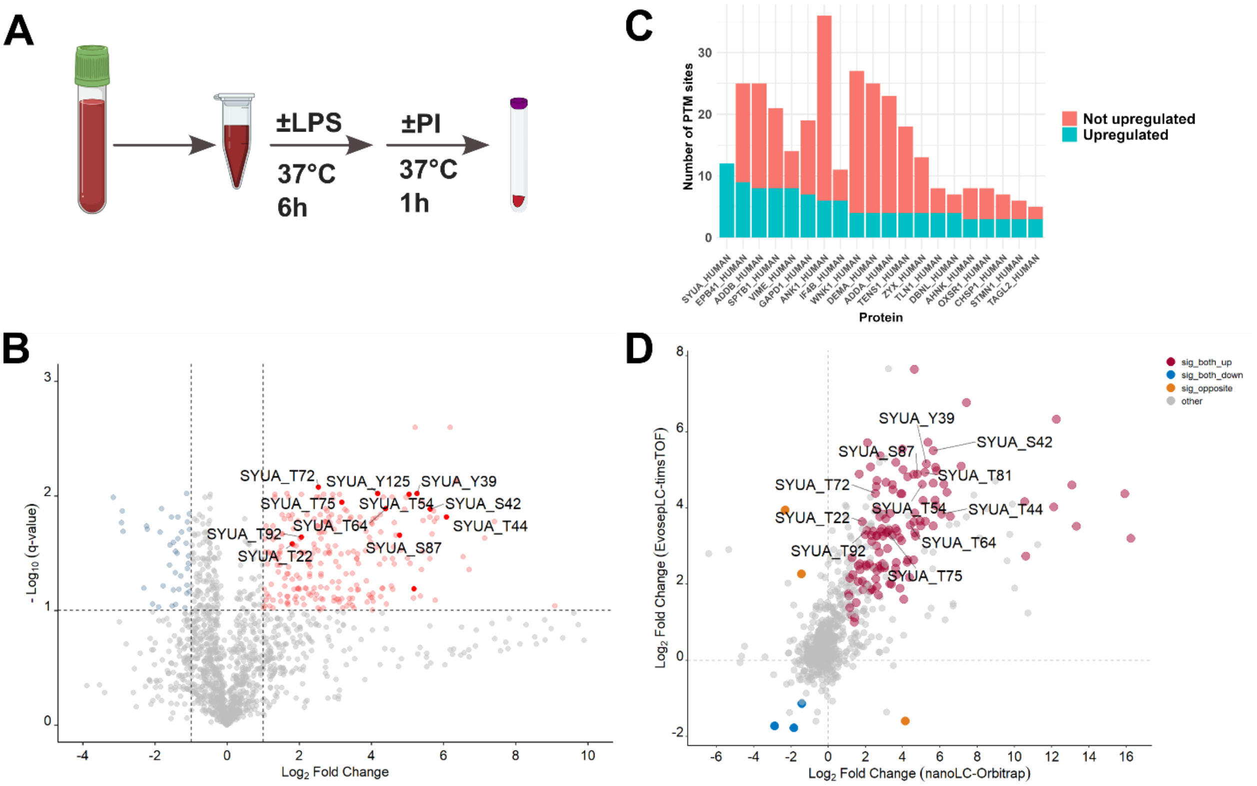


**Figure S4. Effects of phosphatase inhibition on the whole blood phosphoproteome. A**) Heparinized blood from n=4 healthy controls was treated ± 1000 ng/mL *E. coli* LPS at 37 °C for 6h and ± a cocktail of phosphatase inhibitors (PI; 1 mM sodium orthovandate, 500 nM calyculin A and 50 µM deltamethrin) for 1 h followed by loading on Mitra devices, trypsinization and TiO enrichment of phosphopeptides and MS/MS using nanoLC, EvosepLC and SPE-CZE (see Methods). PTM site data was log2-transformed and PI versus sham (no LPS) conditions were analyzed using a paired t-test with Benjamini-Hochberg FDR correction. **B**) A volcano plot of nanoLC-OT-DIA data with alpha synuclein phosphosites highlighted. **C**) A stacked bar plot of nanoLC-OT-DIA data showing the top 20 protein groups with the highest number of significantly upregulated phosphosites and other quantified phosphosites **D**) A correlation plot of differentially abundant phosphosites quantified by nanoLC-OT-DIA versus EvosepLC-timsTOF-DIA.

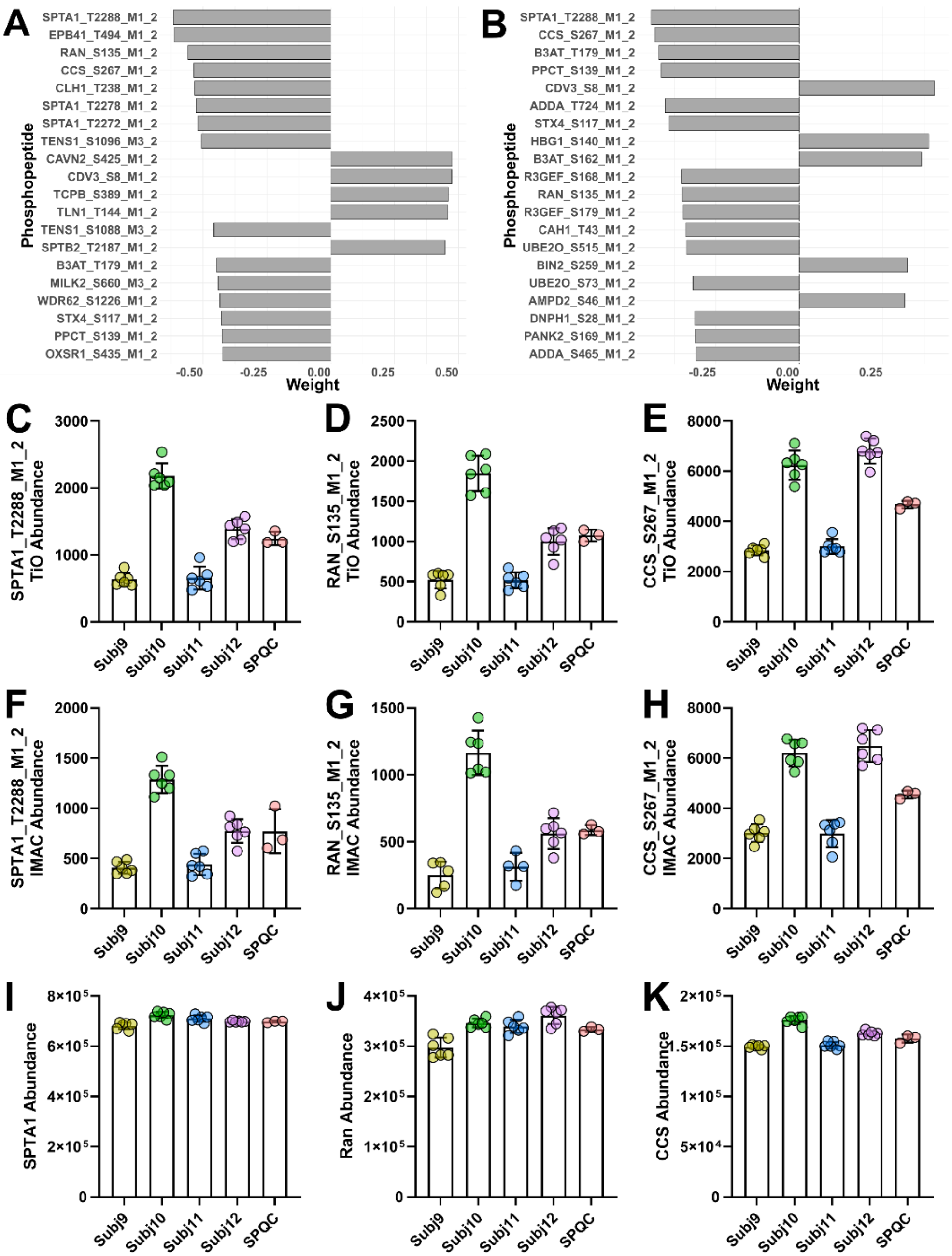


**Figure S5. MOFA2 analysis of phosphosites in wet blood stability study. A-B**) Top 20 phosphosites weighting factor 1 of the wet blood MOFA analysis which used (**A**) TiO or (**B**) Ti-IMAC enrichment of phosphopeptides. **C-H**) Select phosphosites in the top 20 weighting factor 1 that were validated between (**C-E**) TiO enrichment and (**F-H**) Ti-iMAC enrichment versus (**I-J**) total abundance from unenriched proteome.
